# Generation of Infant-dedicated Fine-grained Functional Parcellation Maps of Cerebral Cortex

**DOI:** 10.1101/2021.11.24.469844

**Authors:** Fan Wang, Han Zhang, Zhengwang Wu, Dan Hu, Zhen Zhou, Li Wang, Weili Lin, Gang Li, for UNC/UMN Baby Connectome Project Consortium

## Abstract

Infancy is a dynamic and immensely important period in human brain development. Studies of infant functional development using resting-state fMRI rely on precisely defined cortical parcellation maps. However, available adult-based functional parcellation maps are not applicable for infants due to their substantial differences in functional organizations. Fine-grained infant-dedicated cortical parcellation maps are highly desired but remain scarce, due to difficulties ranging from acquiring to processing of infant brain MRIs. In this study, leveraging 1,064 high-resolution longitudinal rs-fMRIs from 197 infants from birth to 24 months and advanced infant-dedicated processing tools, we create the first set of infant-specific, fine-grained cortical functional parcellation maps. Besides the conventional folding-based cortical registration, we specifically establish the functional correspondences across individuals using functional gradient densities and generate both age-specific and age-common fine-grained parcellation maps. The first set of comprehensive brain functional developmental maps are accordingly derived, and reveals a complex, hitherto unseen multi-peak fluctuation development pattern in temporal variations of gradient density, network sizes, and local efficiency, with more dynamic changes during the first 9 months than other ages. Our proposed method is applicable in generating fine-grained parcellations for the whole lifespan, and our parcellation maps will be available online to advance the neuroimaging field.

## 1 Introduction

The dynamic brain functional development during the first two postnatal years is important for establishing cognitive abilities and behaviors that could last a lifetime (*1–3*). As a prerequisite for understanding how the brain works and develops, cortical parcellation maps provide a repository that helps cortical area localization, network node definition, inter-subject comparison, inter-study communication, and comparison, as well as reducing data complexity while improving statistical sensitivity and power (*4*). In the functional aspect, researchers used to reveal and understand the cortical network topography by clustering cortical vertices into parcels that are different from each other in functional architecture using adult resting-state fMRI (rs-fMRI) data (*5, 6*). Although these clustering-based methods can produce convincing results given a limited number of clusters, they are not suitable for fine-grained parcellations (e.g., ≫ 100 parcels), as they usually result in considerable disjointed fragments that are hardly explainable. To this point, recent adult parcellations (*7–10*) started to use gradient-based methods, i.e., the functional gradient density, to delineate sharp changes of resting-state functional connectivity (RSFC) patterns to promote the meaningfulness and accuracy of parcel boundaries.

All the abovementioned studies derived functional parcellation maps using adult data, which are not suitable for infant studies featuring dynamic brain structural and functional development, due to enormous differences in brain functional organization between infants and adults (*11, 12*). Therefore, infant-specific cortical functional parcellation maps are highly desired, but remain scarce, due to difficulties in both acquiring high-resolution infant brain multi-modal MR images and challenges in processing infant MR images that typically have prominently dynamic imaging appearance and extremely poor tissue contrast (*1, 2, 13*). Of note, another critical issue of using the abovementioned methods for generating infant functional parcellations is that these methods typically computed the functional gradient density map for a cohort directly based on cortical folding-based registration and extensive spatial smoothing of functional connectivity, which, however, cannot lead to accurate functional alignment across individuals, due to large variation between folding and functional areas. Thus many vital details of the functional architecture are blurred and inherently missed in the resulting functional parcellation maps.

In this paper, we aim to generate the first set of infant-specific, high-resolution, fine-grained functional parcellation maps on the cortical surface to significantly accelerate early brain development studies. To this end, this study leverages a large high-resolution dataset with 1,064 rs-fMRI scans and 394 T1-weighted and T2-weighted structural MRI scans from birth to 2 years of age, as part of the UNC/UMN Baby Connectome Project (*14*). To ensure accuracy, all MR images are processed using an extensively validated, advanced infant-dedicated cortical surface-based pipeline (*15*). To establish accurate cortical functional alignment across individuals, we propose a novel method to first compute the functional gradient density map of each infant scan, rather than for the whole cohort in the traditional way, to capture fine-grained functional patterns, and then co-register all functional gradient density maps across individuals based on both cortical folding and functional gradient information. Following steps detailed in Fig. 1, our derived group-average functional gradient density maps capture much more details of cortical functional architecture than the conventional method, thus enabling us to generate fine-grained age-specific cortical parcellation maps of infants at multiple ages, i.e., 3, 6, 9, 12, 18, and 24 months of age. To facilitate infant studies requiring parcel-to-parcel correspondences across ages, we also generate age-common parcellation maps that are suitable for all ages during the first two years. Our infant functional parcellation maps will soon be released to the public to greatly contribute to the pediatric neuroimaging research community.

**Fig. 1.**
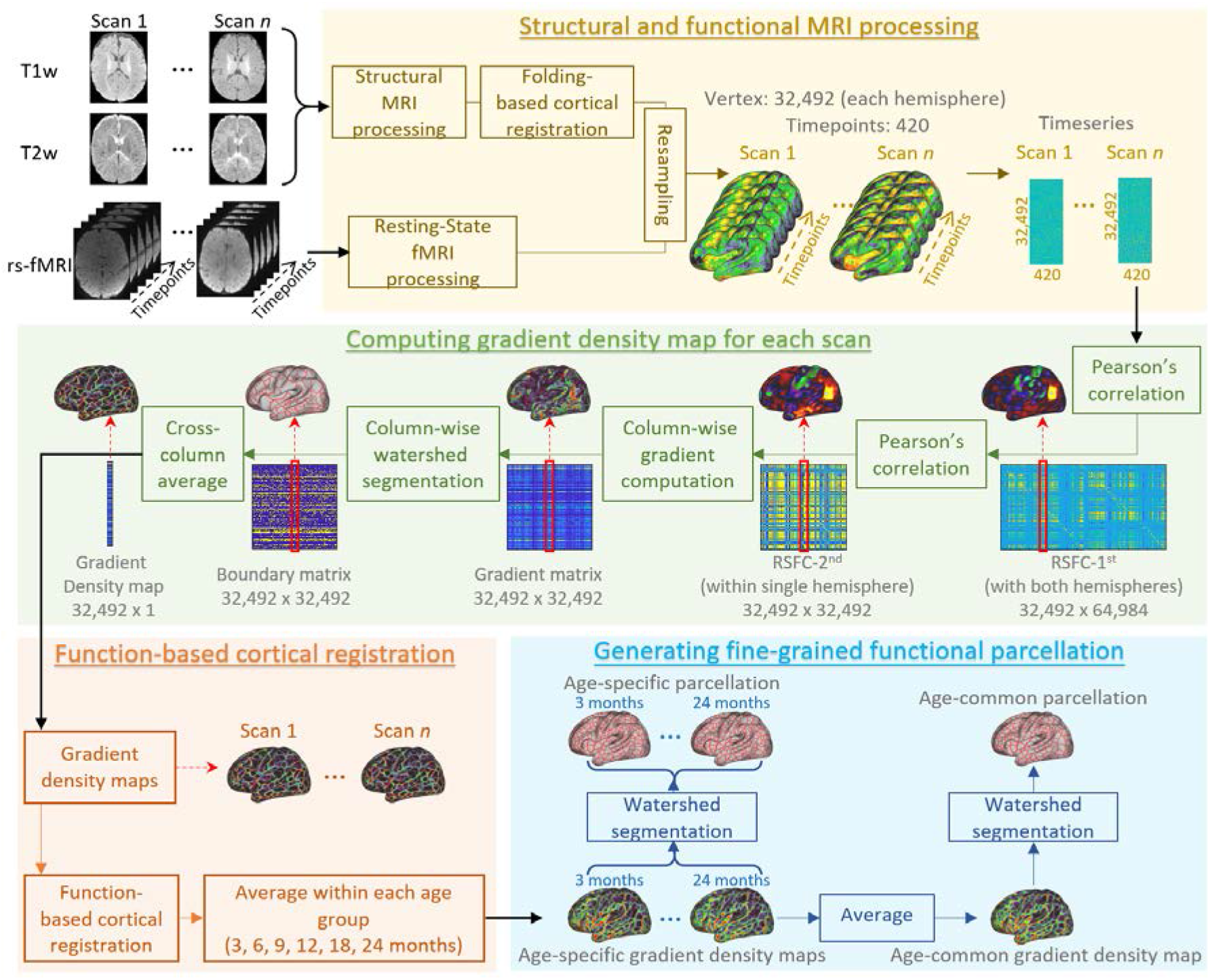
The procedure of infant parcellation using functional gradient density. Our major steps include structural and functional MRI processing, computing gradient density map for each scan, function-based cortical registration, and generating fine-grained functional parcellation.

## 2 Results

We unprecedentedly investigated the fine-grained cortical surface-based functional parcellation maps of the infant cerebral cortex using 1,064 high-resolution (2 × 2 × 2 mm^3^) resting-state fMRI scans from 197 healthy infants, with subject demographics shown in Table 1 and Fig. 8. To capture detailed patterns of sharp transition between cortical areas, after the conventional cortical folding-based inter-individual cortical registration, the gradient density map of cortical functional connectivity was computed on each scan of each individual and further used as a reliable functional feature for function-based registration for establishing functionally more meaningful cortical correspondences across individuals. This resulted in considerably detailed visualization of functional boundaries on the cerebral cortex, which was used to create the infant-dedicated fine-grained cortical functional parcellation maps. Detailed steps of the proposed method are illustrated in Fig. 1.

**Table 1.** Demographic information of each age group from the longitudinal dataset under study.

| Age Group | Age Range (days) | fMRI Scans | AP Scans | PA Scans | Structural MRI Scans (males/females) |
| --- | --- | --- | --- | --- | --- |
| **3M** | 10~144<br>(97.7±35.1) | 109 | 55 | 54 | 52 (27/25) |
| **6M** | 145~223<br>(183.3±23.3) | 172 | 85 | 87 | 56 (24/32) |
| **9M** | 224~318<br>(278.6±24.1) | 139 | 67 | 72 | 54 (27/27) |
| **12M** | 319~410<br>(366.4±21.4) | 151 | 76 | 75 | 57 (24/33) |
| **18M** | 411~591<br>(494.9±51.8) | 244 | 119 | 125 | 91 (40/51) |
| **24M** | 592~874<br>(723.6±67.5) | 249 | 122 | 127 | 84 (42/42) |
| **Total** | 10~874<br>(410.9±219.1) | 1064 | 524 | 540 | 394 (184/210) |

### 2.1 Advantage of the Proposed Method

The functional gradient density maps of 3-month infant scans generated by different methods are compared in Fig. 2, which demonstrates the advantage of our proposed method. Specifically, Fig. 2 (a) shows the group-average gradient density map directly computed using the group-average RSFC-2^nd^ as in (*7, 9*). Fig. 2 (b) shows the group-average gradient density map based on individual gradient density maps, in which we first computed a gradient density map on the RSFC-2^nd^ of each individual and then averaged them across individuals. Fig. 2 (c) shows the group-average gradient density map generated by the proposed method, where all individual gradient density maps are co-registered using the gradient density as a functional feature and then further averaged across individuals. It can be observed that major patterns of the functional gradient density in Fig. 2 (a) are well preserved in Fig. 2 (c) (with some examples pointed out with white arrows), which implies the meaningfulness of the functional gradient density patterns in Fig. 2 (c). Most importantly, Fig. 2 (c) exhibits much more detailed and clear patterns of the functional gradient density, compared to Fig. 2 (a) and (b), especially in the temporo-occipital, parietal, and lateral prefrontal areas, indicating the advantage of performing the 2^nd^ round of co-registration based on functional gradient density. Consequently, the functional gradient density maps generated by the proposed method are able to capture fine-scaled architectures of infant functional connectivity while maintaining the major functional patterns, thus leading to more meaningful fine-grained functional parcellation maps. We also show the gradient density maps of three random subjects in Fig. 2 (d), which further demonstrates that our method can well capture these important and detailed functional gradient patterns, which are usually missed by the compared methods.

**Fig. 2.**
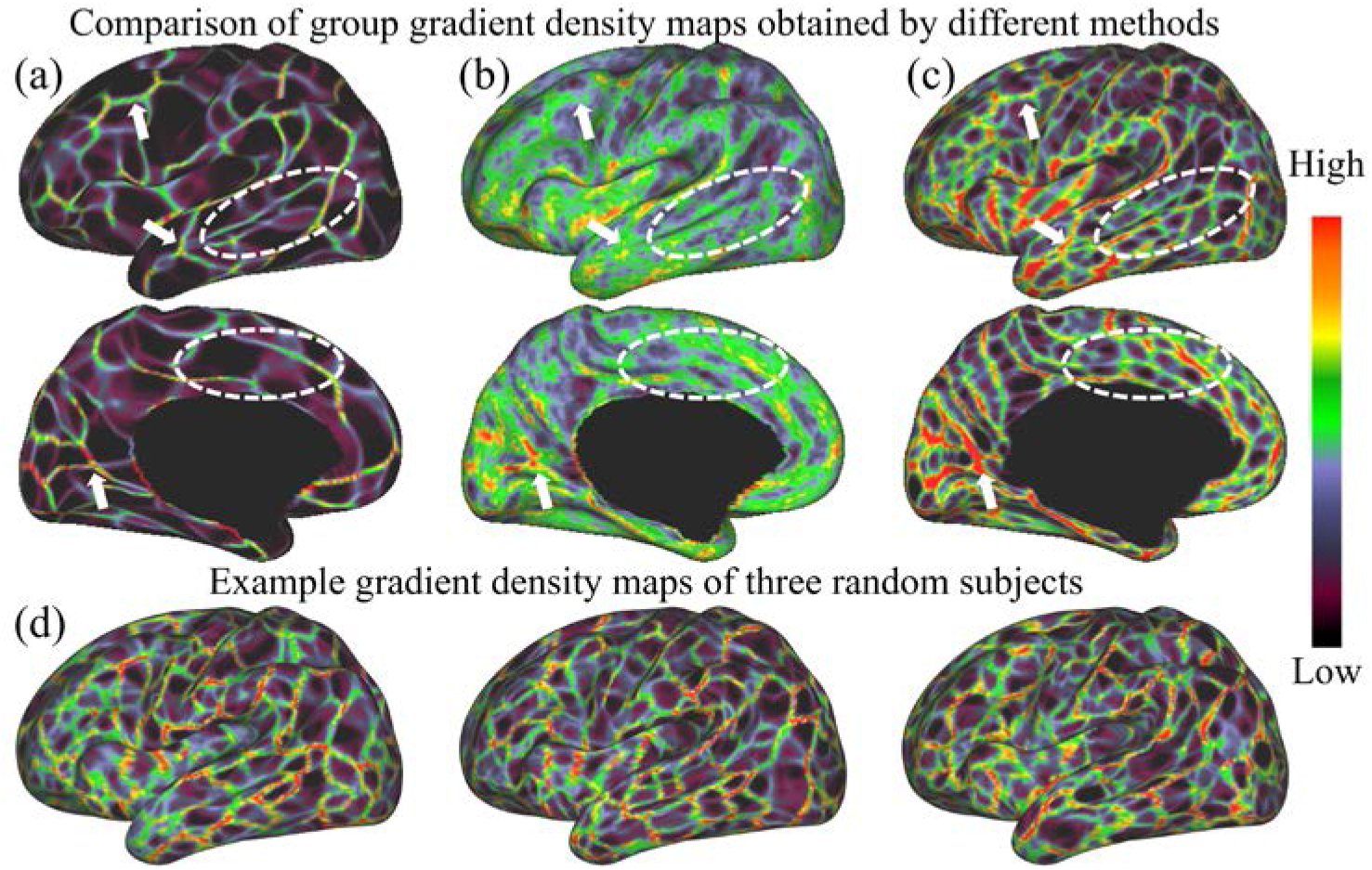
Comparison of the group-average functional gradient density maps on the 3-month age group generated by different methods. (a) The gradient density map computed directly on the population-average connectivity matrix. (b) The gradient density map computed on each individual and then averaged across individuals. (c) The gradient density map generated by our method, which computes the average of individual gradient density maps after co-registration of them based on both cortical folding and functional gradient density. White arrows point out consistent gradient density patterns using different methods, and white dashed circles show some more detailed and fine-grained patterns revealed by our method. (d) Example gradient density maps of three random subjects. This figure demonstrates that some detailed gradient patterns in the individual cortex are usually missed by other methods, and can be well captured by our method.

To test whether the gradient density maps are reproducible, we randomly divided subjects into two non-overlapping parts and computed the dice ratio between two gradient maps after thresholding. A higher dice ratio indicates higher reproducibility. By repeating this experiment 1,000 times, the overall dice ratio reaches 0.9295 ± 0.0021, indicating the high reproducibility of our results.

### 2.2 Age-specific Functional Gradient Density and Parcellation Maps

The age-specific functional gradient density maps are computed by averaging gradient density maps of subjects in corresponding age groups and results at 3, 6, 9, 12, 18, and 24 months are shown in Fig. 3 (a). As can be observed, the major gradient patterns are distributed bilaterally symmetrically on the cortex, like the central sulcus, superior temporal gyrus, middle temporal gyrus, parieto-occipital fissure, and calcarine fissure. Nevertheless, certain gradient patterns exhibit hemispheric differences. For example, the precentral gyrus in the right hemisphere has a higher gradient density than that in the left hemisphere. All these spatial distributions of functional gradient density remain largely consistent across ages.

**Fig. 3.**
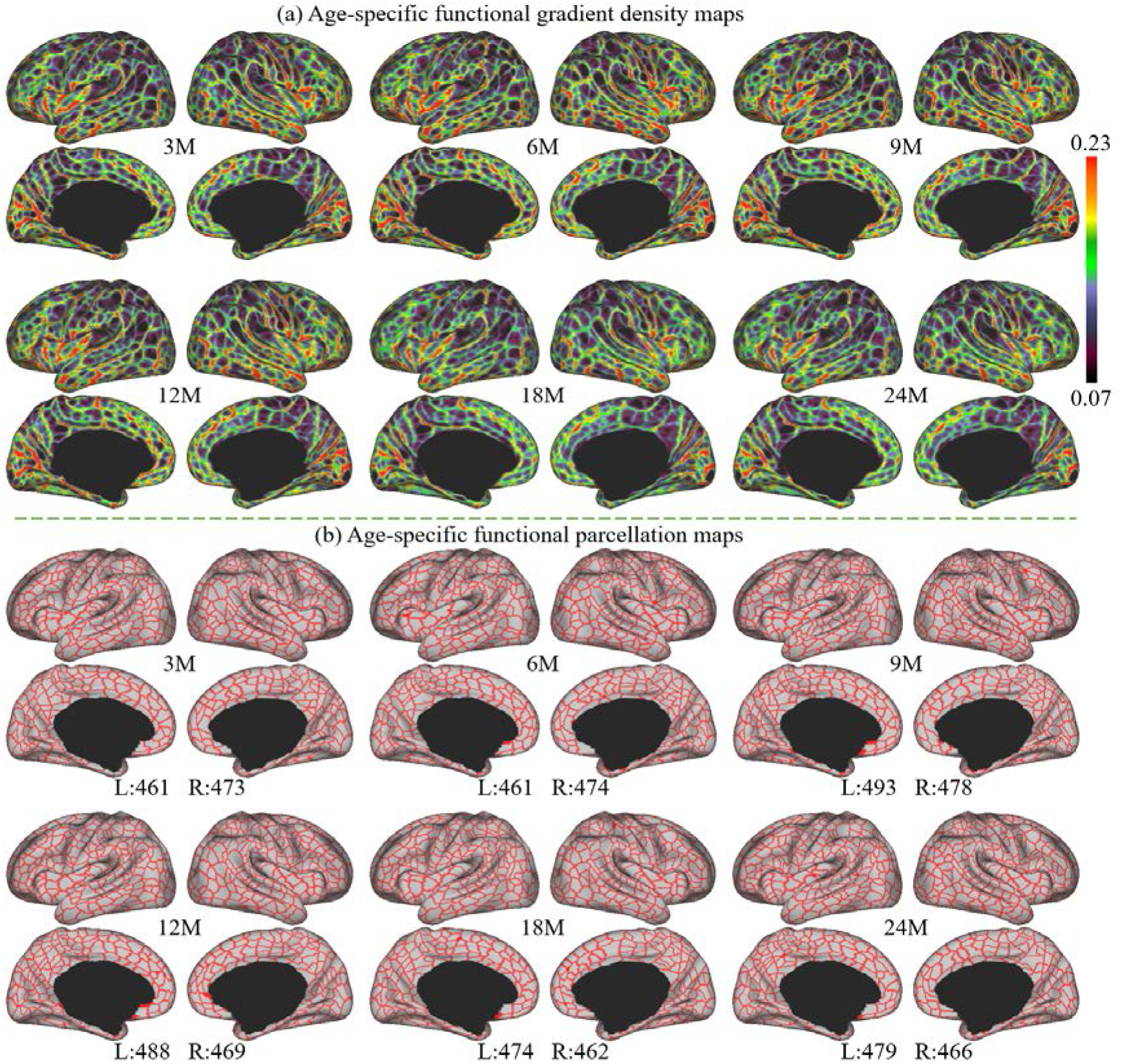
(a) Age-specific functional gradient density maps. (b) Age-specific fine-grained functional parcellation maps, with parcel numbers noted for each age.

Age-specific cortical parcellation maps derived from these functional gradient density maps are presented in Fig. 3 (b). These maps were obtained using a watershed algorithm without thresholding or any manual editing. It can be observed that major gradient density patterns are well reflected as parcellation boundaries. Due to some slight differences in age-specific gradient density maps, the resulting age-specific parcellation maps show different parcel numbers. However, all parcel numbers fall between 461 to 493 parcels per hemisphere, and parcel numbers show slight changes that follow a multi-peak fluctuation, with inflection ages of 9 and 18 months of age.

To evaluate the consistency of gradient density across different age groups, we thresholded and binarized the age-specific functional gradient density maps to their top 50% and 25% gradient density. These binary maps were summed up, resulting in a gradient density overlap map indicating its age consistency shown in Fig. 4 (b). In these maps, “one” stands for high gradient densities that appeared in only one age group, and “six” represents high gradient densities that appeared in all six age groups. It is worth noting that most high gradient densities are repeatedly detected in all six age groups, suggesting the high consistency of majorities of high gradient densities in all age groups.

**Fig. 4.**
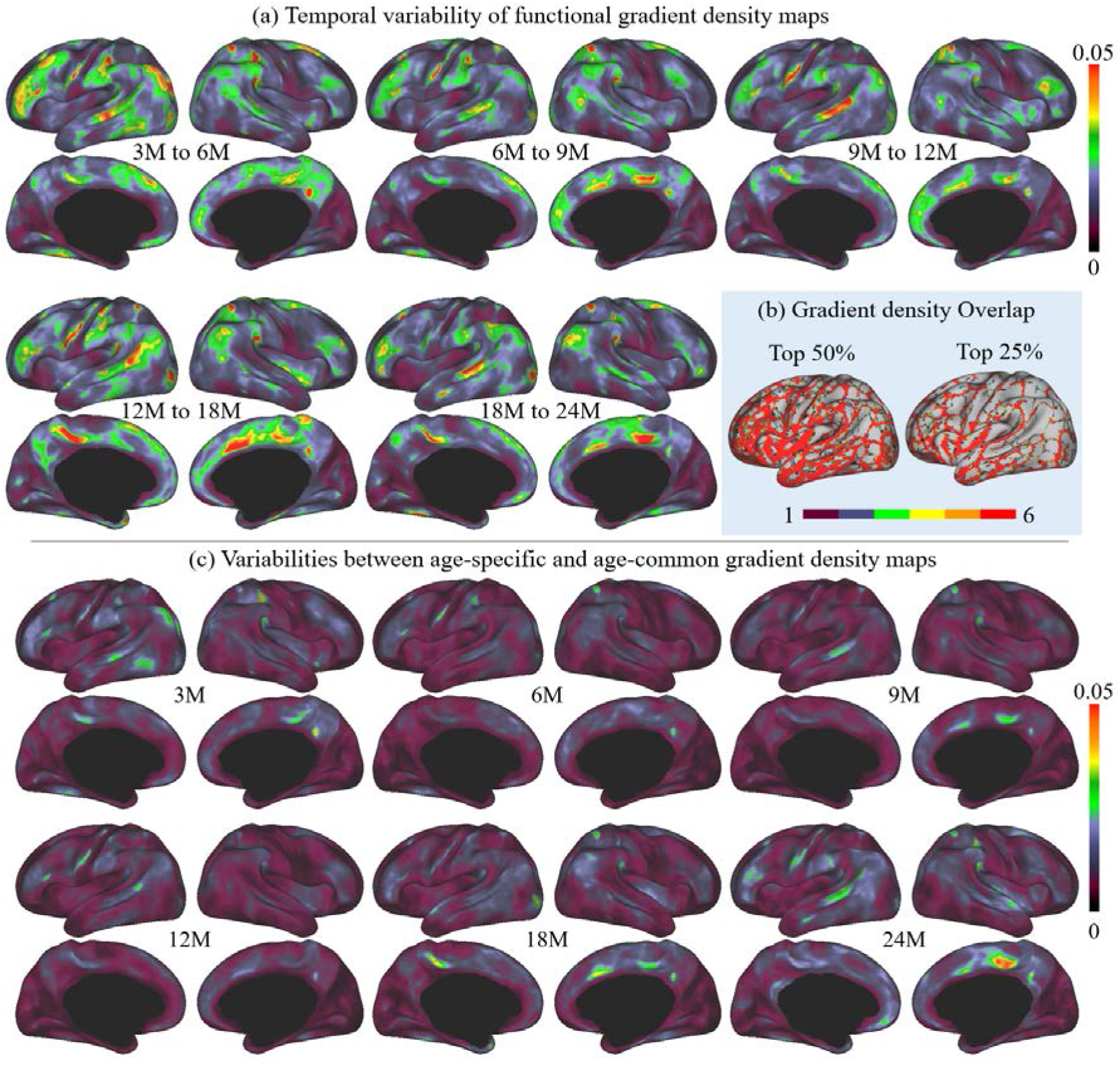
(a) Temporal variabilities of functional gradient density maps between every two consecutive ages. (b) Consistency of high gradient density across ages. (c) Variabilities between each age-specific functional gradient density map and that of the age-common map.

Further, to better illustrate the functional architecture development, we computed the temporal variability of gradient density maps between neighboring age groups, as shown in Fig. 4 (a). In general, the temporal variabilities of functional gradient density are at a relatively low level (<=0.05) at all age intervals. Across all ages, high temporal variabilities are mainly presented in high-order association areas, including the left middle and inferior frontal, middle temporal, right superior frontal, precuneus medial prefrontal, and bilateral supramarginal, posterior superior temporal, and medial frontal areas. Other regions mostly exhibit low temporal variabilities, especially in the sensorimotor and medial occipital regions. Keeping this spatial distribution, the temporal variability shows a multi-peak fluctuation, where the gradient density decreases from 3-6 to 6-9 months, followed by an increase during 9-12 and 12-18 months, and drops again during 18-24 months.

### 2.3 Age-common Functional Gradient Density and Parcellation Maps

Since infant functional MRI studies typically involve multiple age groups, it is highly desired to have an age-common functional parcellation map that features parcel-to-parcel correspondences across ages, so that it can be conveniently employed for all ages during infancy. Therefore, we also computed the age-common gradient density map (Fig. 5 (a)) as the average of the functional gradient density maps of all six age groups. The variabilities between the age-common gradient density map and each age-specific gradient density map are illustrated in Fig. 4 (c). Compared to the temporal variability between neighboring age groups (Fig. 4 (a)), the age-common gradient density map shows small variability to all age-specific maps. The spatial distributions of high and low variabilities remain mostly similar to that of the temporal variabilities between neighboring age groups, with high variability presented in some high-order association cortices and low variability in unimodal cortices. Consequently, it can be speculated that the age-common gradient density map can be used to generate an age-common parcellation map that is suitable for all subjects from birth to 2 years of age.

**Fig. 5.**
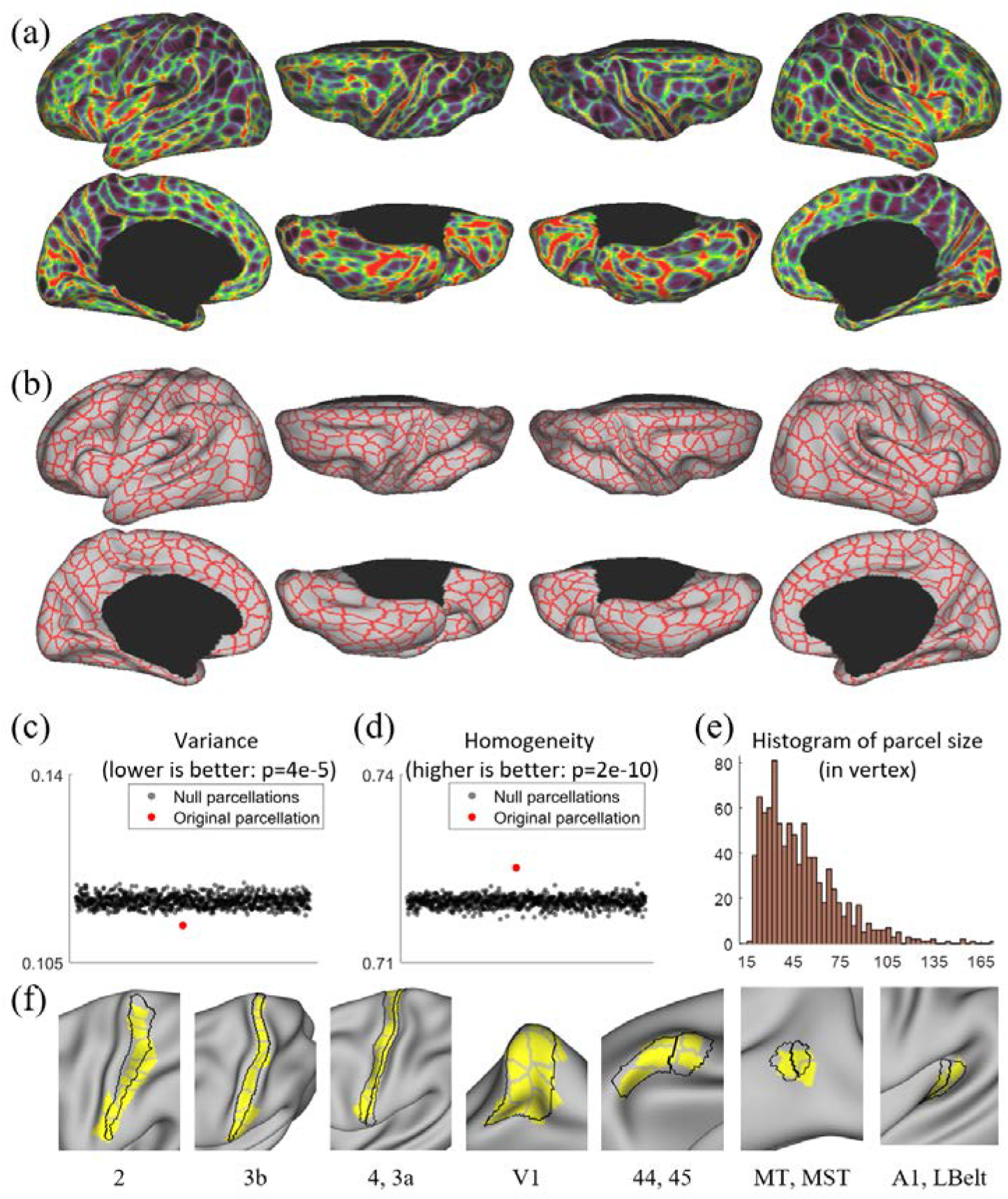
(a) Age-common gradient density map. (b) Age-common parcellation map (864 parcels, L: 432, R: 432). (c) Our age-common parcellation shows significantly lower variance compared to the null parcellations. (d) Our age-common parcellation shows significantly higher homogeneity compared to null parcellations. (e) The histogram of parcel size, where parcel sizes are counted in vertices. (f) Some parcels correspond to known cortical areas defined by multi-modal features in adults (*4*).

The resulted age-common functional parcellation map based on the age-common gradient density map is shown in Fig. 5 (b), which has 864 parcels in total (L: 432, R: 432) excluding the medial wall. It should be noted that we manually removed some apparently over-segmented regions after using the watershed algorithm, and prior to that, we had 903 parcels in total (L: 448, R: 455). The parcel boundaries of the age-common parcellation map are well aligned with high gradient density regions and show largely bilaterally symmetric patterns of the areal organization. In the following development-related analyses in this study, we mainly employed the age-common parcellation map to facilitate comparisons of infants across ages.

Compared to existing fine-grained parcellation maps, such as the multi-modal adult parcellation (*4*), the age-common infant parcellation map has comparatively smaller and more evenly distributed parcel sizes and shapes. Also, as shown in Fig. 5 (f), some areas of our parcels show substantial overlap with the known cortical areas of adults, such as the visual areas V1, MT, MST, sensorimotor areas 2, 3, 4, auditory areas A1, LBelt, and language areas 44, 45. To further examine the validity of our parcellation map, we compared it with 1,000 null parcellation maps in terms of variance and homogeneity, with the results shown in Fig. 5 (c) and (d). It can be observed that our parcellation map shows significantly higher homogeneity (*p*=2e-10) and lower variance (p=4e-05), indicating the meaningfulness of the resulting parcellation map.

### 2.4 Network Organization and Development

We performed network clustering of the generated parcels in each age group to reveal the early development of functional network organization. The number of networks for each age group is determined separately according to the random split-half stability analysis. Empirically, the network number is set as 2 to 30, and the stability plots are shown in Fig. 6 (a). Higher stability suggests a better clustering result, hence a more meaningful network organization. Overall, when the network number surpasses 15, the stability does not show a substantial raise or decrease, indicating the network numbers likely hold less than 15. Hence, we look for the cluster number on a ‘peak’ or prior to a ‘descending cliff’, which guarantees high stability or more significant network numbers. As a result, we find that the most suitable cluster numbers for different age groups are 7 networks for 3 months, 9 networks for 6 months, 10 networks for 9, 12, 18, and 24 months. Of note, we choose10 networks for 18 months so as to be consistent during development, even though it is neither a peak nor a cliff.

**Fig. 6.**
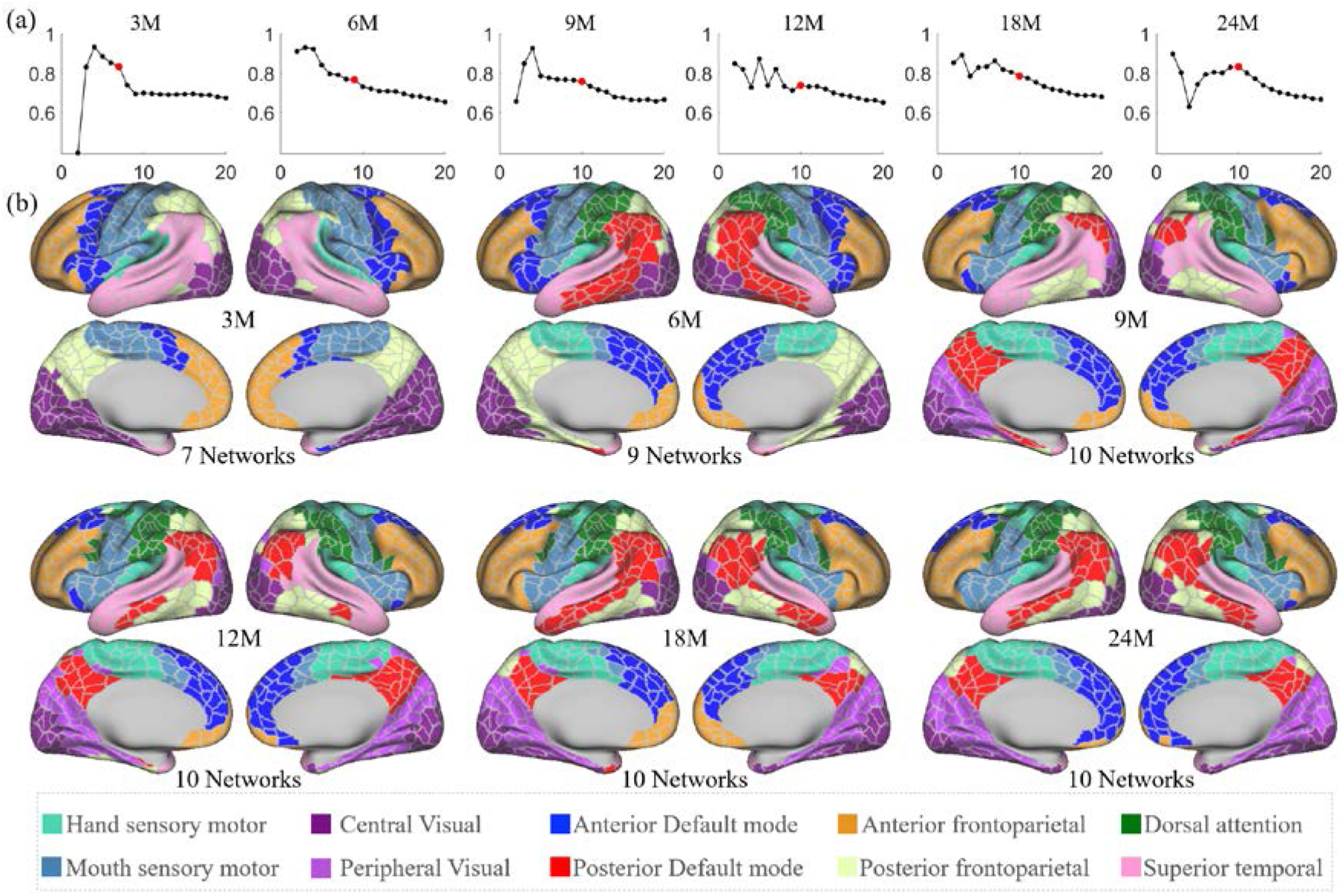
(a) Stabilities of different network numbers of different age groups computed by repeating 200 times random split-half test. The selected numbers are highlighted in solid red dots. (b) Discovered functional network organization of parcels during infantile brain development, color-coded by corresponding networks denoted below.

The spatiotemporal patterns of the discovered functional network organization are shown in Fig. 6 (b). Overall, changes in network structure from 3 to 9 months are more extensive than those from 9 to 24 months. Specifically, the sensorimotor network splits into two subnetworks from 3 to 6 months, and the boundary between them moved toward the ventral direction from 6 to 9 months. The hand sensorimotor expands, while the mouth sensorimotor shrinks and both stabilize after 9 months. The auditory network is distinguished at 3 months and merges into the hand sensorimotor at 6 months. The visual network splits into peripheral and central visual subnetworks from 6 to 9 months and remains stable until a slight shrinkage at 24 months.

Other networks exhibit more complex development with multi-peak fluctuation of the size in certain networks. Specifically, the anterior default mode network expands from 3 to 6 months, and shrinks from 6 to 9 months and from 12 to 18 months, and expands thereafter. The lateral posterior default mode network that emerged at 6 months shrinks from 6 to 9 months and then expands from 9 to 18 months; while the medial posterior default mode network emerged at 9 months only lightly shrinks thereafter. The anterior and posterior default mode networks develop to the adult-like pattern at 18 months, while till 24 months, they are still detected as two separate networks. The superior temporal network shrinks from 3 to 6 months, expands from 6 to 9 months, and then shrinks again from 9 to 24 months. The anterior frontoparietal network gradually shrinks from 3 to 24 months, except for an expansion from 12 to 18 months. On the medial surface, the posterior frontoparietal network expands to include the parahippocampal gyrus from 3 to 6 months and then disappears by 9 months. On the lateral surface, the posterior frontoparietal network expands from 3 to 9 months to include the inferior temporal part and becomes stable thereafter. The dorsal attention network is seen at 6 months and evolves to the adult-like pattern at 9 months and keeps stable thereafter.

### 2.5 Parcel-wise Development

Homogeneity of functional connectivity can be used as a criterion for characterizing functional development. Fig. 7 (a) shows the parcel-wise homogeneity development during infancy. Our results suggest that the overall parcel-wise homogeneity shows a monotonic decrease trend during the first two years by maintaining similar relative spatial distribution. Higher homogeneities are located in the sensorimotor, paracentral, posterior insula, inferior parietal, posterior superior temporal, lateral occipital, and occipital pole. Low homogeneities are presented in lateral prefrontal, medial frontal, anterior insula, inferior temporal, and temporal pole.

**Fig. 7.**
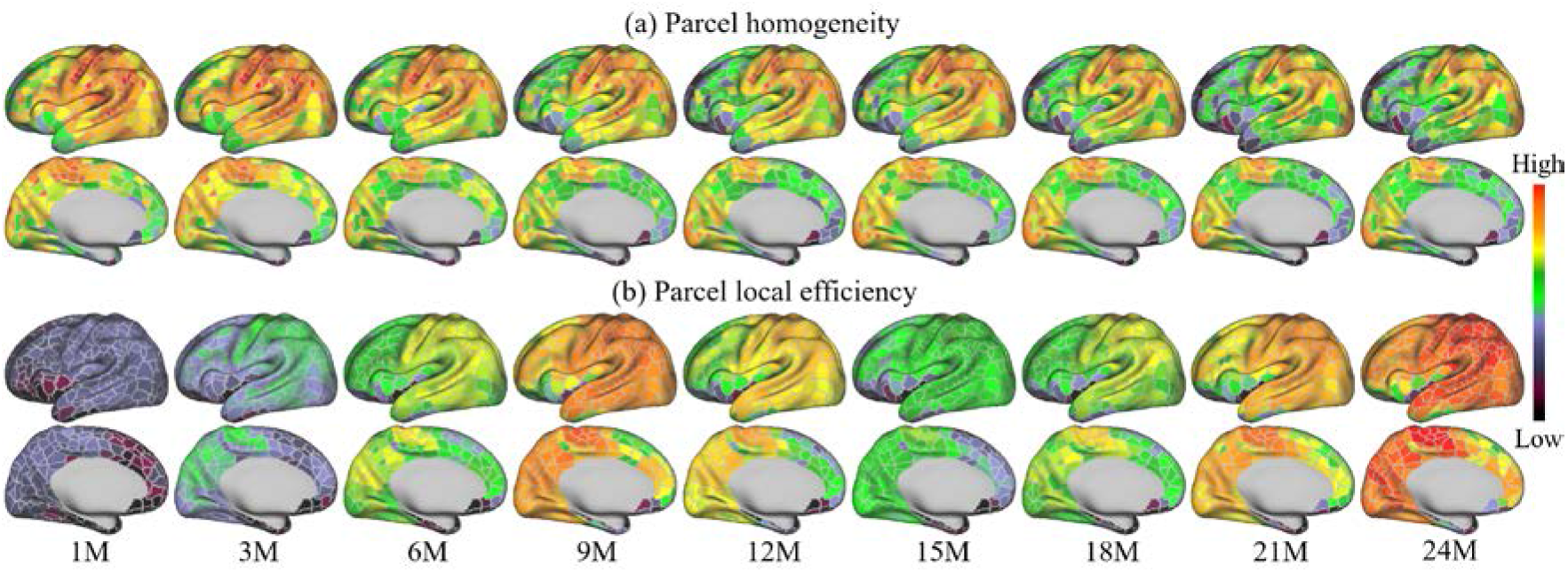
Development of parcel homogeneity and local efficiency during infancy.

Fig. 7 (b) shows the development of the local efficiency of each parcel. Overall, the local efficiency also exhibits a strong multi-peak fluctuation, with inflection ages observed at 9 and 15 months. Parcels with low efficiency are located in the lateral superior frontal, medial superior frontal, orbitofrontal, ventral insula, and anterior inferior temporal cortices. Parcels with high local efficiency are mainly observed in sensorimotor, paracentral, parietal, and precuneus regions.

## 3 Discussion

In this study, we created the first set of both age-specific and age-common, infant-dedicated, fine-grained, and cortical surface-based functional parcellation maps using functional gradient density maps. We analyzed the spatiotemporal patterns of age-specific functional gradient density maps and found that age-common functional gradient density maps are suitable for creating fine-grained functional parcellation maps for all ages in the infant cohort. We validated the meaningfulness of the parcellation and showed that its boundaries substantially reproduced known areal boundaries, and its parcels featured high homogeneity and low variance. Finally, we illustrated the infantile development in network structure, parcel homogeneity, and parcel local efficiency.

This study used the functional gradient density as a feature for improving functional alignment across individuals, in addition to the conventional cortical folding features used by previous adult functional parcellations (*7, 9*). As a result, our method not only captured important coarse gradient patterns discovered by previous methods, but also revealed much more detailed areal boundaries at a remarkable resolution, as compared in Fig. 2. The main reason is that previous studies solely relied on cortical folding-based registration, thus inevitably suffered from significant inter-subject variability in the relation between cortical folding and functional areas, leading to less accurate inter-subject functional correspondences. For infant-dedicated functional parcellations, the only one available is the volumetric-based parcellation based on image registration and clustering generated by Shi et al. (*16*), without any advanced surface-based processing and registration. In contrast, our parcellation maps are generated based on the cortical surface, which well respects the topology of the convoluted cerebral cortex, and avoids mixing signals from opposite sulcal banks and different tissues, leading to more accurate functional signals resampling, smoothing, computation and registration. Moreover, our parcellations leveraged high-quality 2 mm isotropic fMRI data that densely covers the first two years, instead of data with a coarse resolution of 4 mm isotropic centering at birth, 1 and 2 years of age (*16*).

Standing upon the detailed gradient density patterns by the proposed method, we generated age-specific fine-grained parcellation maps for 3, 6, 9, 12, 18, and 24 months of age. We found that the temporal variability (temporal changes) of the functional gradient density generally decreases during most age intervals, except for a slight increase from 9 to 12 months. This may suggest that the development of functional architecture gradually slows down during the first two years. We also found that high temporal variabilities mostly presented in high-order association cortices, implying that they are developing at a more considerable pace compared to unimodal cortices. Besides the age-specific parcellations, we also generated an age-common parcellation that suites infants at all ages to help brain development-related studies under two considerations. First, infant studies typically involve subjects of different ages, and it is not convenient to use different parcellations for comparison between different ages. Second, our age-common gradient density map shows low variability to all age-specific gradient density maps and therefore can generate the representative fine-grained functional parcellation map that is suitable for all ages during infancy. However, we will make both the age-specific and age-common functional parcellation maps accessible to the public in the case that some researchers may still prefer age-specific parcellation maps.

This study successfully augmented the resolution of the existing cortical parcellations from ~300 to ~900 areas, which represents a finer architecture of brain functional organizations compared to previous ones. This fine-grained cortical organization is also in line with Eickhoff et al. (*17*), where they believe that 200-300 areas are not the ultimate resolution for cortical parcellations due to the multi-hierarchical formation of the brain. Glasser et al. (*4*) also consider 360 as a lower bound for cortical parcellations since each parcel can be represented as a combination of several smaller regions. Consequently, our fine-grained infant cortical parcellation maps provide a great platform for analyzing pediatric neuroimaging data with a greatly boosted resolution, thus leading to more meaningful discoveries on the fine-scaled functional architecture of infant brains.

Besides, it is worth noting that our parcellation increased the resolution in a meaningful way. First, our functional gradient density maps are highly reproducible. By separating subjects into non-overlapping parts, their gradient density patterns are repeated with a dice ratio of ~0.93. Second, our age-common infant parcellation shows high accordance in some specific cortical areas defined by Glasser et al. (*4*), which is recognized as the state-of-the-art adult parcellation map. As illustrated in Fig. 5 (d), our gradient density map-derived parcellation contains parcels that have substantial overlap with the known adult area V1 defined by Glasser et al. (*4*). Other known cortical areas of adults, such as sensorimotor areas 2, 3, and 4, were also overlapped with a combination of several parcels in our parcellation. These observations that parcel borders conform to some adult cortical areas lend substantial visual validity to the parcellation.

When applying parcellation as a tool to explore infant brain functional development, our results reveal complex multi-peak fluctuations in several aspects, including parcel number, temporal variation of gradient density, network organization, and local efficiency. To the best of our knowledge, this complex fluctuation development trend is not reported in previous literature and should fill an important knowledge gap for infantile brain functional development. These functional developmental patterns are very different from early brain structure development, where the cortical thickness follows an inverted-“U” shaped trajectory, while the surface area and cortical volume monotonically increase following a logistic curve. The multi-peak fluctuations potentially mirror different milestones of behavioral/cognitive abilities, which likely emerge at different ages during infancy (*30*). However, the underlying mechanisms of such developmental patterns remain to be further investigated.

For network organization (Fig. 6 (b)), at 3 months, networks likely groups vertices with close spatial locations, resulting in networks being more dependent on the local anatomy. After 9 months, the primary functional systems reach steady and present adult-like patterns, while high-order functional networks still show substantial differences compared to the adult-like pattern. Our results suggest that a primitive form of brain functional networks is present at 3 months, which is largely consistent with recent studies suggesting that most resting-state networks are already in place at term birth (*18–20*). Besides, our results also suggest that, compared to high-order functional networks, the primary functional system is more developed in infants. This confirms previous findings in infant cortical thickness development (*21*), suggesting that the primary functional systems develop earlier than high-order systems.

At the network level, the sensorimotor system splits into two sub-networks, i.e., the mouth- and hand-sensorimotor at 6 months, which were also observed in infants and toddlers (*22*). The visual network is split into central (primary) visual and peripheral (high-order) visual cortices at 9 months and well maintains this pattern until 24 months. This subdivision of mouth- and hand-sensorimotor networks is also found in adults (*5*). The high-order functional systems, including the default mode, frontoparietal, and dorsal attention network, exhibit considerable development during 3 to 9 months, followed by some minor adjustments from 12 to 24 months. A previous infant study (*19*) also demonstrated that functional network development shows more considerable change in the first year compared to the second year. At 24 months, both default mode and frontoparietal networks show a lack of strong cross-lobe connections. Though several studies identified some prototypes of cross-lobe connection (*19, 23*), their links seem not as strong as to be stably distinguished (*24–26*). Our results suggest that the high-order functional networks are far more from established at 24 months of age. It is worth noting that, the size changes of networks can be quite subtle between a short time interval, which emphasizes the importance of using a fine-grained parcellation map.

The parcel homogeneity measures the development within parcels. Our result shows (Fig. 7 (a)) that unimodal cortices, including the sensorimotor, auditory, and visual areas, show high homogeneity, which is largely consistent with adults (*7*). However, the inferior parietal and posterior superior temporal cortices, which show high homogeneity in infants, are observed low homogeneity in adults (*7*). Besides, the prefrontal area, which shows relatively low homogeneity in infants, seems to develop to a medium-to-high homogeneity in adults. Almost all parcels are observed decreased homogeneity with age. This is likely related to the development of brain function, especially in high-order cortices, which show increased heterogeneity, and consequently decreased homogeneity. Among the high-order association cortices, the prefrontal area has the lowest homogeneity, followed by temporal and then parietal regions, suggesting different levels of functional development.

Local efficiency measures a different developmental aspect – it represents the connection of parcels to neighbors. Higher local efficiency is usually related to higher functional segregation (*11*). Our results (Fig. 7 (b)) suggest that local efficiency shares certain similar spatial distribution with homogeneity – they both increase in anterior to posterior and ventral to dorsal directions. During development, the local efficiency shows a complex developmental trend: although 24 months shows a strong increase compared to 1 month, there is a dip from 12 to 21 months that should be noted. The age-related increase of local efficiency was previously found from 18 months to 18 years (*27*), 5 to 18 years (*28*), and 12 to 30 years (*29*), and is likely explained by progressive white matter maturation (*27*). This trajectory of local efficiency is not contradictory to the previous studies (*31, 32*), since they only measured the averaged local efficiency of all nodes to reflect network characterization, thus missing important characteristics of parcel-level local efficiency. This further stresses the importance of performing parcel-wise analyses and the significance of fine-grained infant cortical parcellations.

## 4 Conclusion

In summary, for the first time, this study constructed a comprehensive set of cortical surface-based infant-dedicated fine-grained functional parcellation maps. To this end, we developed a novel method for establishing functionally more accurate inter-subject cortical correspondences. We delineated age-specific parcellation maps at 3, 6, 9, 12, 18, and 24 months of age as well as an age- common parcellation map to facilitate studies involving infants at different ages. Our parcellation maps were demonstrated meaningful by comparing with known areal boundaries and through quantitative evaluation of homogeneity and variance of functional connectivity. Leveraging our infant parcellation, we provide the first comprehensive visualizations of the infant brain functional developmental maps on the cortex and reveal a complex multi-peak fluctuation functional development trend, which will serves as valuable references for future early brain developmental studies. Our generated fine-grained infant cortical functional parcellation maps will be released to the public soon to greatly advance pediatric neuroimaging studies.

## 5 Methods

### 5.1 Subjects and Image Acquisition

Subjects in this study are from the UNC/UMN Baby Connectome Project (BCP) data90set (*14*). The BCP focuses on normal early brain development, where all infants were born at the gestational age of 37-42 weeks and free of any major pregnancy and delivery complications. In this study, 394 high-resolution longitudinal structural MRI scans were acquired from 197 (90 males and 107 females) typically developing infants, as demonstrated in Fig. 8. Images were acquired on 3T Siemens Prisma MRI scanners using a 32-channel head coil during natural sleeping. T1-weighted images (208 sagittal slices) were obtained by using the three-dimensional magnetization-prepared rapid gradient echo (MPRAGE) sequence: TR (repetition time)/TE (echo time)/TI (inversion time) = 2,400/2.24/1,600 ms, FA (flip angle) = 8º, and resolution = 0.8×0.8×0.8 mm^3^. T2-weighted images (208 sagittal slices) were acquired with turbo spin-echo sequences (turbo factor = 314, echo train length = 1,166 ms): TR/TE = 3,200/564 ms, and resolution = 0.8×0.8×0.8 mm^3^ with a variable flip angle. All structural MRI data were assessed visually for excessive motion, insufficient coverage, and/or ghosting to ensure sufficient image quality for processing.

**Fig. 8.**
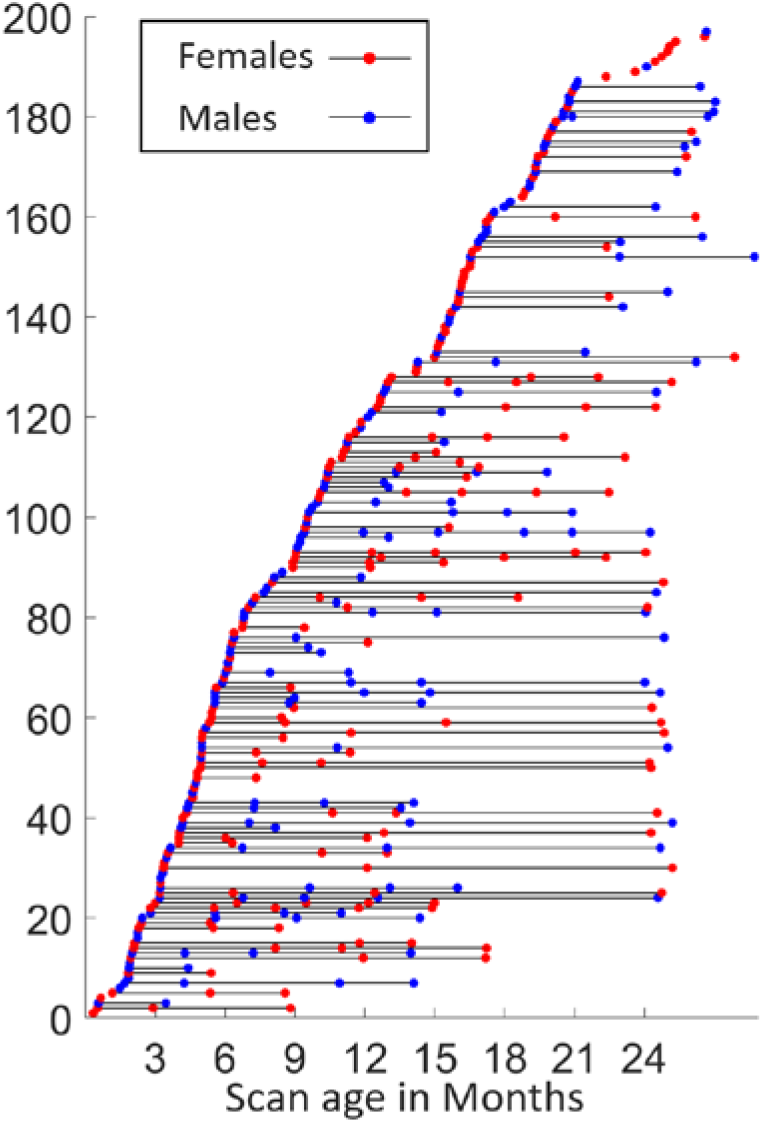
Longitudinal distribution of scans. Each point represents a scan with its scanned age (in months) shown in the x-axis, with males in blue and females in red, and each horizontal line represents one subject, with males in blue and females in red.

For the same cohort, 1,064 high-resolution resting-state fMRI (rs-fMRI) scans were also acquired using a blood oxygenation level-dependent (BOLD) contrast sensitive gradient echo echo-planar sequence: TR = 800 ms, TE = 37 ms, flip angle = 80°, field of view (FOV) = 208×208 mm, 72 axial slices per volume, resolution = 2×2×2 mm^3^, total volumes = 420 (5 min 47 s). The rs-fMRI scans include 524 anterior to posterior (AP) scans and 540 posterior to anterior (PA) scans, which are two opposite phase-encoding directions for better correction of geometric distortions.

### 5.2 Structural MRI Processing

All T1-weighted and T2-weighted MR images were processed using an infant-specific pipeline detailed in (*15, 33*), which have been extensively validated in many infant studies (*21, 34–41*). The processing procedure includes the following main steps: 1) Rigid alignment of each T2-weighted image onto its corresponding T1-weighted image using FLIRT in FSL (Smith et al., 2004); 2) Skull stripping by a deep learning-based method (*42*), followed by manual editing to ensure the clean skull and dura removal; 3) Removal of both cerebellum and brain stem by registration with an atlas; 4) Correction of intensity inhomogeneity using the N3 method (*43*); 5) Longitudinally-consistent segmentation of brain images as white matter (WM), gray matter (GM), and cerebrospinal fluid (CSF) using an infant-dedicated deep learning-based method (*44*); and 6) Separation of each brain into left and right hemispheres and filling non-cortical structures.

### 5.3 Resting-State fMRI Processing

Infant rs-fMRI processing was conducted according to an infant-specific functional pipeline (*31, 45, 46*). The head motion was corrected using FSL, as well as the spatial distortions due to gradient non-linearity. The rs-fMRI scans were then registered to the T1-weighted structural MRI of the same subject using a boundary-based registration approach (*47*). All of the transformations and deformation fields were combined and used to resample the rs-fMRI data in the native space through a one-time resampling strategy. After conservative high-pass filtering with a sigma of 1,000 s to remove linear trends in the data, individual independent component analysis was conducted to decompose each of the preprocessed rs-fMRI data into 150 components using MELODIC in FSL. An automatic deep learning-based noise-related component identification algorithm was used to identify and remove non-signal components to clean the rs-fMRI data (*48*).

### 5.4 Cortical Surface Reconstruction and Mapping

Based on the tissue segmentation results, inner, middle and outer cortical surfaces of each hemisphere of each MRI scan were reconstructed and represented by triangular meshes with correct topology and accurate geometry, by using a topology-preserving deformable surface method (*33, 49*). Before cortical surface reconstruction, topology correction on the whiter matter surface was performed to ensure the spherical topology of each surface (*50*). After surface reconstruction, the inner cortical surface, which has vertex-to-vertex correspondences with the middle and outer cortical surfaces, was further smoothed, inflated, and mapped onto a standard sphere (*51*).

To ensure the accuracy in longitudinal analysis during infancy, it is necessary to perform longitudinally-consistent cortical surface registration (*15*). Specifically, 1) for each subject, we first co-registered the longitudinal cortical surfaces using Spherical Demons (*52*) based on cortical folding-based features, i.e., average convexity and mean curvature. 2) Longitudinal cortical attribute maps were then averaged to obtain the intra-subject mean surface maps. 3) For each hemisphere, all intra-subject mean surface maps were then co-registered and averaged to get the population-mean surface maps. 4) The population-mean surface maps were aligned to the HCP 32k_LR space through registration to the “fsaverage” space as in (*53*). By concatenating the three deformation fields of steps 1, 3 and 4, we directly warped all cortical surfaces from individual scan spaces to the HCP 32k_LR space. These surfaces were further resampled as surface meshes with 32,492 vertices, thus establishing vertex-to-vertex correspondences across individuals and ages. All results were visually inspected to ensure sufficient quality for subsequent analysis. The inner and outer cortical surfaces were used as a constraint to resample the rs-fMRI time courses onto the middle cortical surface with 32,492 vertices using the HCP workbench (*54*), and the time courses were further spatially smoothed on the middle cortical surface with a small Gaussian kernel (σ = 2.55 *mm*).

### 5.5 Generation of Fine-grained Cortical Functional Parcellation Maps

In this section, we describe detailed steps for generating fine-scaled infant cortical functional parcellation maps (see Fig. 1). Specifically, we first describe the computation of the gradient density map of functional connectivity for each scan, followed by a function-based registration step based on gradient density maps. Then, we detail the computation of both “age-specific” and “age-common” parcellation maps based on the functional registration results and our evaluation scheme. At last, we introduce how we use the parcellation maps to discover the functional network organization development, as well as parcel homogeneity and local efficiency during infancy.

#### Computation of Individual Functional Gradient Density Map

The gradient density of functional connectivity (*7*) identifies sharp changes of RSFC, thus intrinsically representing the transition from one functional parcel to another, and is widely used in generating meaningful fMRI-based cortical parcellations in adult studies (*7–9*). For each fMRI scan of each infant subject, the computation of the gradient density of functional connectivity on the cortical surface is summarized in the following steps. 1) For each fMRI scan, the functional connectivity matrix is built by pair-wise correlating each vertex with all other cortical vertices in the CIFTI file to create a 32k×64k RSFC matrix for each hemisphere. 2) Each RSFC matrix is transformed to z scores using Fisher’s r-to-z transformation. 3) For each fMRI scan, the z-transformed RSFC of each vertex is correlated with all cortical vertices within the same hemisphere, creating a 2^nd^ order correlation matrix (RSFC-2^nd^) sized 32k×32k for each hemisphere. 4) For some scan visits consisting of both AP and PA scans, all RSFC-2^nd^ matrices of the same visit from the same subject are averaged, so that all subjects contribute equally, even they may have different numbers of scans in one visit. 5) The gradient of functional connectivity is computed on the RSFC-2^nd^ as in (*4*), resulting in a 32k×32k gradient matrix per hemisphere. 6) By performing the watershed-based boundary detection (*7*) on the gradient matrix, we obtain 32k binary boundary maps per hemisphere. 7) The functional gradient density map is defined as the average of 32k binary boundary maps.

#### Cortical Surface Registration based on Functional Gradient Density

Previous studies mostly computed population-based functional gradient density map, where cortical surfaces were usually co-registered to a common space using only cortical folding-based features. However, due to the highly-variable relationship between cortical folds and functions, especially in high-order association regions, researchers recently are getting more aware of the necessity of functional features-based registration (*55, 56*). To this end, in addition to cortical folding-based co-registration, we further use the gradient density of functional connectivity as a meaningful functional feature to perform a second-round of co-registration of cortical surfaces for the purpose of more accurate functional alignment.

Specifically, based on cortical folding-based surface co-registration, 1) the functional gradient density maps of all scans are averaged to generate the population-mean functional gradient density map. 2) To improve inter-individual cortical functional correspondences, the functional gradient density map of each scan is then aligned onto the current population-mean functional gradient density map using Spherical Demons (*52*) by incorporating functional gradient density as a feature.

3) All warped functional gradient maps are then resampled and averaged to obtain the newly improved population-mean functional gradient density map with sharper and more detailed functional architecture. 4) Steps 2 and 3 are repeated iteratively until no visually observed changes in the population-mean functional gradient density map (4 iterations in our experiment). After this procedure, all individual functional gradient density maps are co-registrated, thus establishing functionally more accurate cortical correspondences across individuals.

#### Generation of Parcellation Maps based on Functional Gradient Density

##### Age-specific Parcellation Maps

To capture the spatiotemporal changes of fine-grained cortical functional maps during infancy, we group all scans into 6 representative age groups, i.e., 3, 6, 9, 12, 18, and 24 months of age based on the distribution of scan ages. For each age group, we compute the age-specific group-average functional gradient density maps by averaging the gradient density maps of all scans within the group, without any smoothing. Detailed information of each age group is reported in **Table 1**. A watershed method is then applied on each age-specific group-average functional gradient density map to generate the corresponding functional parcellation maps (*7*). This watershed segmentation algorithm starts by detecting local minima in 3-ring neighborhoods, and iteratively grows the region until reaching ambiguous locations, where vertices can be assigned to multiple regions. These locations appear to be borders that separate parcels and reflect putative boundaries of functional connectivity according to the functional gradient density maps.

##### Age-common Parcellation Maps

Ideally, the age-specific parcellation maps are the more appropriate representation of the cortical functional architecture at the concerned age. However, many neuroimaging studies involve infants across multiple ages, thus the age-specific parcellation maps may not be proper choices due to different parcel numbers and variation in parcel boundaries across ages, thus inducing difficulties in across-age comparisons. To facilitate infant studies involving multiple age groups, we also compute an age-common gradient density map, which is the average of all 6 age-specific functional gradient density maps without any smoothing, so that each age group contributes equally to the age-common map. According to the age-common functional gradient density map, we generate the age-common functional parcellation map using the watershed segmentation method as well. The subsequent parcellation evaluation, functional network architecture and longitudinal development analyses are performed using the age-common parcellation maps.

### 5.6 Evaluation of Parcellation Maps

#### Reproducibility

Ideally, a functional gradient density-based parcellation map should extract robust common gradient information that shows the transition between parcels. We thus test if the gradient density map is reproducible on different subjects. Therefore, randomly divided “generating” and “repeating” groups (*7, 9*) are used to calculate mean gradient density map, separately. These two maps are then binarized by keeping only 25% highest gradient density as in (*7, 8*), and the dice ratio overlapping index between the two binarized maps is calculated to evaluate the reproducibility of the functional gradient map. This process is repeated multiple times (1,000 times in this study) to get a reliable estimation.

#### Homogeneity

The functional gradient density-based parcellation identifies large gradients, representing sharp transition in functional connectivity pattern and avoiding large gradients inside parcels as much as possible. Meanwhile, a parcel that accurately represents a cortical area should not only be distinct from its neighbors in functional connectivity pattern, but also has a homogenous functional connectivity pattern across all vertices inside. Therefore, we estimate the homogeneity of each parcel as in (*7*). Specifically, we first compute the mean correlation profile of each vertex across all subjects. Next, the correlation patterns of all vertices within one parcel is entered into a principal component analysis; the percentage of the variance that is explained by the largest principal component is used to represent the homogeneity of this parcel.

#### Variance

As the functional connectivity pattern within a parcel should be relatively uniform, we also measure the variability of the connectivity pattern within each parcel, with smaller variability indicating greater uniformity and hence higher parcellation quality. Specifically, for each parcel, we first obtain a matrix with each column representing subject-average z-score of functional connectivity profile of one vertex in the parcel. Then we compute the sum of standard deviation of each row to represent the variability of this parcel. The average variability of all parcels is used to represent the variability of the parcellation map.

As parcellation maps usually have different numbers, sizes and shapes in parcels, to have fair comparison and be consistent with (*7, 8*), we compare our parcellation maps with ‘null parcellations’. The null parcellations are generated by rotating by a random amount along x, y and z axes on the 32,492 spherical surfaces, which relocate each parcel while keeping the same number and size of parcels. We compare both variability and homogeneity of our parcellation and that of the random rotated null parcellations. Notably, in any random rotation, some parcels will inevitably be rotated into the medial wall, where no functional data exist. The homogeneity/variance of a parcel rotated into the medial wall is not calculated; instead, we assign this parcel the average homogeneity/variance of all random versions of the parcel that were rotated into non-medial-wall cortical regions.

#### Variability Between *Functional Gradient Density* Maps

A variability map visualizes the variability or dissimilarity between two functional gradient density maps, and is estimated as follows. For a vertex *ν*, a surface patch centering at *ν* is extracted (10-ring neighborhood in this study), and two vectors ***p***_1_ and ***p***_2_ within this patch are then extracted from two functional gradient density maps. Their variability at *ν* is computed as 0.5 × (1 − *corr*(***p***_1_, ***p***_2_)), where *corr*(⋅,⋅) stands for Pearson’s correlation. As a result, the variability/dissimilarity is within the range of [0, 1], where high value stands for high variability/dissimilarity and vice versa. In this study, we mainly measure the variability between functional gradient density maps in two aspects: 1) the temporal variability, which computes the variability of functional gradient density maps between two consecutive age groups to reflect the developmental changes of the gradient density maps; 2) the variability between the age-common functional gradient density map and each age-specific functional gradient density map, for quantitatively evaluating whether it is appropriate to use the age-common parcellation maps for all 6 age groups.

### 5.7 Functional Development Analysis

#### Functional Network Detection

To discover the developmental evolution of large-scale cortical functional networks, we employ a network discovery method (*5*) to each of the 6 age groups. Specifically, for each subject in each age group, given *n* parcels, we first compute the average time course of each parcel (excluding the medial wall), and compute the correlation of the average time courses between any two parcels. This results in a *n* × *n* matrix, which is further binarized by setting the top 10% of the correlations to one and the rest to zero. For each age group, all *n* × *n* matrices are averaged across individuals independently. A clustering algorithm (*57*) is then applied to estimate networks of parcels with similar connectivity profiles.

To determine the optimal cluster number *k* for each age group, we employ the random split-half test to compute the stability for each *k*, with higher stability corresponding to more meaningful clustering results. Specifically, for each age group, we randomly split all subjects into two folds and run the clustering algorithm separately to obtain two independent clustering results *c*_1_ and *c*_2_, and the similarity between *c*_1_ and *c*_2_ is evaluated using the Amari-type distance (*58*). This experiment is repeated 200 times for each age group, and the resulted similarities are averaged to represent the stability for *k*. During this process, the range of *k* is set to [2, 30] according to the existing literature of functional network discovery (*5, 6*).

#### Parcel-wise Development

We computed the homogeneity and local efficiency of each parcel in the age-common parcellation to characterize infantile parcel-wise developmental patterns regarding functional homogeneity and functional segregation, respectively. The homogeneity is computed as described in Section 3.4 for each subject, where higher parcel homogeneity indicates more unified connectivity pattern within the parcel. The local efficiency is computed using the GRETNA Toolkit (*59*) for each subject. Herein, multiple thresholds are used, keeping 50% to 5% connections with 1% as a step, and the area under curve (AUC) is calculated to represent the local efficiency to avoid the influence of connectivity densities. The local efficiency corresponds to the mean information transfer efficiency between a particular parcel and all its connected nodes, which is proportional to the clustering coefficient. Parcels with higher local efficiency can more effectively share information to its connected parcels, and thus help build effective segregated networks. To have intuitive and spatiotemporally detailed views of their development, we use the sliding window technique to compute homogeneity and local efficiency in each age window by averaging all scans within the same age window. The windows are centered at each month, with a window width of 90 days (±45 days) at 2 months of age, increasing 4 days in width for each following month and reaching 182 days (±91 days) at 2 years of age.

## Acknowledgment

This work was partially supported by NIH grants (MH116225, MH117943, MH104324, MH109773, MH123202, MH127544). This work also utilizes approaches developed by an NIH grant (1U01MH110274) and the efforts of the UNC/UMN Baby Connectome Project Consortium.

